# Distribution, Epidemics dynamics and physiological races of wheat stem rust (*Puccinia graminis* f.sp. *tritici* Eriks and E. Henn) on irrigated wheat in the Awash River basin of Ethiopia

**DOI:** 10.1101/2021.03.22.436408

**Authors:** Nurhussein Seid Yesuf, Sileshi Getahun, Shiferaw Hassen, Yoseph Alemayehu, Kitessa Gutu Danu, Zemedkun Alemu, Tsegaab Tesfaye, Netsanet Bacha Hei, Gerald Blasch

## Abstract

*Wheat is one of the high value important major crops of the globe. However, wheat stem rust is considered one of the determinant threats to wheat production in Ethiopia and the globe. So this study was conducted with the objective to assess disease intensity, seasonal distribution dynamics, and genetic variability and to determine virulence spectrum of stem rust in the irrigated wheat areas of Ethiopia*. A total of 137 wheat farms were *evaluated from 2014/15 - 2019/20 in six districts of Awash River basin. Farm plots were assessed every 5 - 10 km interval with ‘X’ fashion, and data on disease incidence, severity, and healthy plants were scored with diseased wheat plant samples collection for stem rust race analysis. The seasonal trend of wheat stem rust disease was also compared to see the future importance of the diseases. The result revealed that the prevalence, incidence, and severity of stem rust were significantly varied among the different districts and seasons in the two regions. The survey results also indicated that about 71.7% of the wheat fields were affected by stem rust during the 2018/19 growing season. The overall incidence and mean severity of the disease during the same season were 49.02% and 29.27%, respectively. During 2019/20 about 63.7% of the wheat fields were affected by stem rust, which, however, the incidence* (*30.97%*) *and severity* (*17.22%*) *were lower than the previous season. Although the seasonal disease distribution was decreased, its spatial distribution was expanding into Lower Awash. The physiological and the genetic race analysis identified four dominant races* (*TTTTF, TKTTF TKKTF, and TTKTF*) *during 2018/19 and additional race* (*TKPTF*) *during 2019/20. Thus races are highly virulent and affect most of the Sr genes except Sr – 31 and Sr – 24. TTTTF and TKKTF are the widest virulence spectrum which affects 90% of the Sr genes. Thus, it can be concluded that the spatial and seasonal distribution of the disease was expanding. Moreover, most of the races were similar with rain-fed production, and thus care must be given for effective management of the diseases to both agro-ecologies. Therefore, these findings provide inputs or insight for breeders to think about the breeding programs in their crossing lines and wheat producers to reduce the damage of the disease in the irrigated ecologies*.

**ETHICAL STATEMENT:** Thus, surveys were conducted with the lateral aim of rust epidemics early warning and monitoring support program in the Awash River basin. Samples for this study were collected from farmers’ fields of the irrigated production areas in the Awash River basin. The disease was an air-borne disease that is difficult to contain. Still, we give maximum care during surveying through spore-free through self-sanitation after Pgt infested field observation to minimize induced disease dissemination to the communities in the production areas that no specific permissions were required for these locations. Field sites are on public access, and *P. graminis* f. sp. tritici is already an air-born pathogen that doesn’t need special protection kinds. This work was our study experience in the endeavor in irrigated wheat technology dissemination.

## 1. INTRODUCTION

Wheat (*Triticum aestivum* L.) is one of the leading cereal grains in the world where more than one-third of the population uses it as a staple food (Kumar *et al*., 2011). The fastest world Population growth is expected to reach nine billion by 2050 (Edmeades *et al*., 2010) and there should be an urgent need to produce highly productive crops like wheat to feed the world population soon (Weigand, 2011). Sub-Sahara Africa (SSA) wheat is produced with a total of 7.5 MT on a total area of 2.9 M ha accounting for 40 and 1.4% of the wheat production in Africa and at global levels, respectively (FAO, 2017). The most important wheat-producing countries in SSA are Ethiopia, South Africa, Sudan, Kenya, Tanzania, Nigeria, Zimbabwe, and Zambia in descending order. Ethiopia accounts for the largest production area (1.7 Mha) followed by South Africa (0.5 Mha) (Wuletaw *et al*., 2018). Wheat production in Ethiopia is expanding at 3.69% from the period of 1979/80-2017/18 (Diriba, 2018). The total annual production of wheat in Ethiopia is 4.64 million tons from 1.7 million hectares in 2018 growing season (CSA, 2019).

Previously, wheat has been predominantly produced in southeastern, central, and northwestern regions of Ethiopia (Tesemma and Belay, 1991) and currently, the Ethiopian government started irrigated wheat production projects as a mechanism to substitute its import. Irrigated crop productivity is generally found to be higher than rain-fed crop productivity, and for decades the Ethiopian government’s policies have promoted irrigation expansion particularly for cereal crops as a method for improving agricultural growth, smoothing production risk, and alleviating rural poverty. The production of wheat in the irrigated agroecology is not only an attractive business but it is a multi-social responsibility to support the country in food self-sufficiency, source of animal feed for pastoralists, and mechanisms to fill the gap for foreign exchange. However, its production and productivity are very low compared with the world average (3.3 t/ha) this could be due to several factors of which biotic (diseases, insects, and weeds), abiotic stress, and low adoption of new agricultural technologies are the major ones (Zegeye *et al*., 2001; Admassu *et al*., 2009). Moreover, Rosegrant and Agcaoili (2010) and Wuletaw *et al*. (2018) also reported that the increasing demand for wheat at the global level, on the one hand, and the challenges facing its production such as climate change, increased cost of inputs, increased intensity of abiotic (drought, heat) and biotic (diseases and pests) stresses, on the other hand, make the wheat demand-supply chain very volatile and at times lead to social unrest.

Among the biotic stresses, wheat stems rust which is caused by *Puccinia graminis* f. sp. *tritici* is caused 100% yield losses (during epidemic years on susceptible cultivar (Park *et al*., 2007; Hailu *et al*., 2015). The surveillance result revealed the predominance of stem rust in East and southern Africa (Hodson *et al*., 2012). In Ethiopia, a lot of hot spot areas for the appearance of virulent genetic diversity of stem rust races were also reported by Hailu *et al*. (2015).

Ethiopian highlands are considered as a hot spot area for the development of stem rust diversity and it becomes a threat when it is supported with a mono-cropping system, continuous-release, and commonality parentage effect on extensive cultivation of CIMMYT originated genotypes Leppik, (1970). Efforts have been done so far by using effective Sr genes in combination with other genes with gene pyramiding and they found great importance as the additive effects of several genes offer the cultivar a wider base stem rust resistance (Abebe *et al*., 2012).

Ethiopia’s prospect of wheat self-sufficiency in the coming two years will be possible with two favorable and realistic cases in addition to positive promising results (MOA, 2020). Increment of production in the rain-fed areas and expansion of production to irrigable lowlands was a strategy to expand production in the country (MOA, 2020). Recent evidence revealed that irrigation wheat was a key factor in increasing wheat production and productivity. The expansion of those production technologies across the main river basins of the country was started from the technological support of Werer Agricultural research center in the Awash River basin. In 2018 about 3502 ha, 2019 about 15,100 ha of land in Afar (Amibara, Dubti, and Gewane), Oromia region (Sire and Jeju), South Omo (Arba-Minch zuria, South Omo), and Somalia (Gode) was targeted (MOA, 2020). The government has a plan to area expansion up to 300,000 ha for 2020 years in Oromia, Amhara Somalia, Southern Nations and Nationalities and Peoples Regional State, and Afar.

The major abiotic production constraints in irrigated lowland areas of Ethiopia are environmental stresses like extreme temperature, soil salinity, drought and flood, soil pH, and salinity (Sileshi *et al*., 2015). However, the stem rust disease was the most important prevalent and severe disease which majorly reduces wheat production in the two regions of the study area except Dubti district in the Afar region; likely the menace could be prominent biotic stress in Ethiopia. Stem rust which also designated as Pgt hereafter has been the most devastating rust disease of all wheat rusts in the country (Admassu *et al*., 2012) and could be the only destructive rust disease in the irrigated lowlands of Ethiopia with environmental factors related to the other two rust pathogens in the disease triangle.

The irrigated lowlands will help to create a suitable micro-climate for the growth of *Pg*t with the ability to change its genetic makeup into new races which will increase the difficulty to discover resistant and high-yielding varieties. Ethiopia is considered to be one of the hotspots for new race development and thus, the new production areas will intensify this theory. Pgt could be effectively controlled through growing resistant varieties knowing the prevalent races were the required knowledge in the variety development (Hei, 2018). *Pgt* prevalence, incidence, and severity parameters with their defined dominant virulent and virulent races give essential information on the gene-to-gene concept to be considered in the breeding programs (Park *et al*., 2011). A huge Pgt outbreak by race TKTTF in 2013 caused up to 100% yield losses in some fields (Sanders, 2011).

Our research hypothesis was to address the questions how was the distributed in irrigated wheat production of Ethiopia? How the disease intensity was going in the irrigated wheat production areas of the Awash River basin? What are the likely national scale economic losses from the *Pgt* based up on the field data set? What are the common race genetic and physiological races of wheat stem rust on irrigated wheat in the Awash River basin of Ethiopia usually anew race expected from new area cultivation, is there a new race? Generate information on the Pgt epidemics dynamics with in recurrent expanding production scenarios. The present study was, therefore, targeted to know the distribution of wheat stem rust prevalence, incidence, and severity and their virulence spectrum to ensure sustainable irrigated wheat production in Awash River basin and similar lowland area irrigated wheat production agro ecological settings (344-1266 m.a.s.l.).

The objective of this study was to determine the diseases intensity distribution, epidemics dynamics, and to analyze the Pgt races in the irrigated belts of Awash River basin.

## 2. MATERIALS AND METHODS

### 2.1. Study Site Description

The field survey was conducted on upper, middle, and lower Awash River basins during 2014/15 to 2019/20 irrigated wheat growing seasons. The study includes seven districts having four (Amibara, Dubti, Samorovia Gellalo, and Gewane) from Afar and three from Oromia regional state (Fentale, Jeju, and Sire). The districts are located in the range between 039° 39’ 20” E and 41°43’40” E and 08° 25’ 50” N and 11°32’20” N (Figure 1).

**Figure 1.**
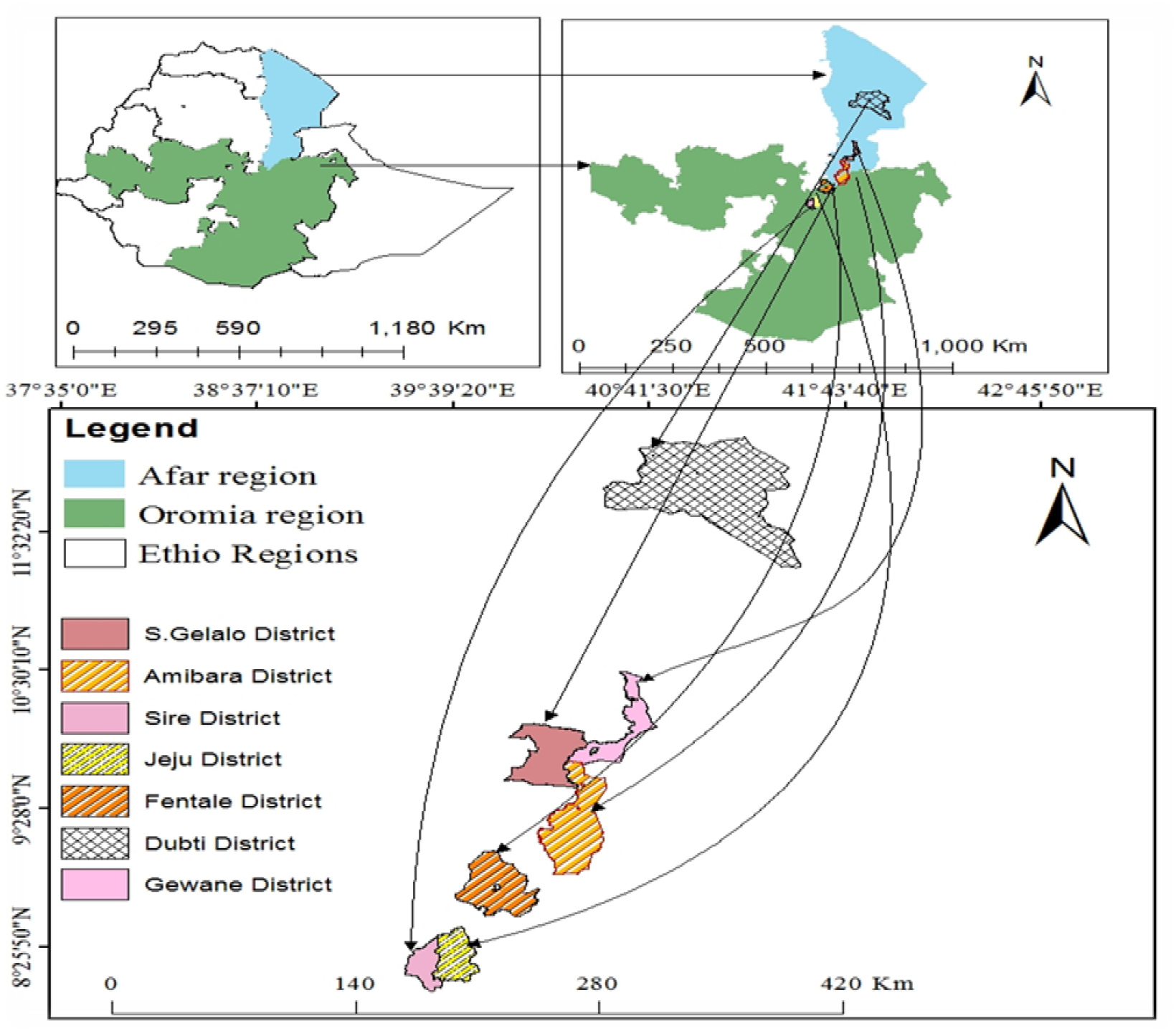
The irrigated wheat production belt status in Awash River basin in 2018/19 cool-season cropping season

**Figure 2.**
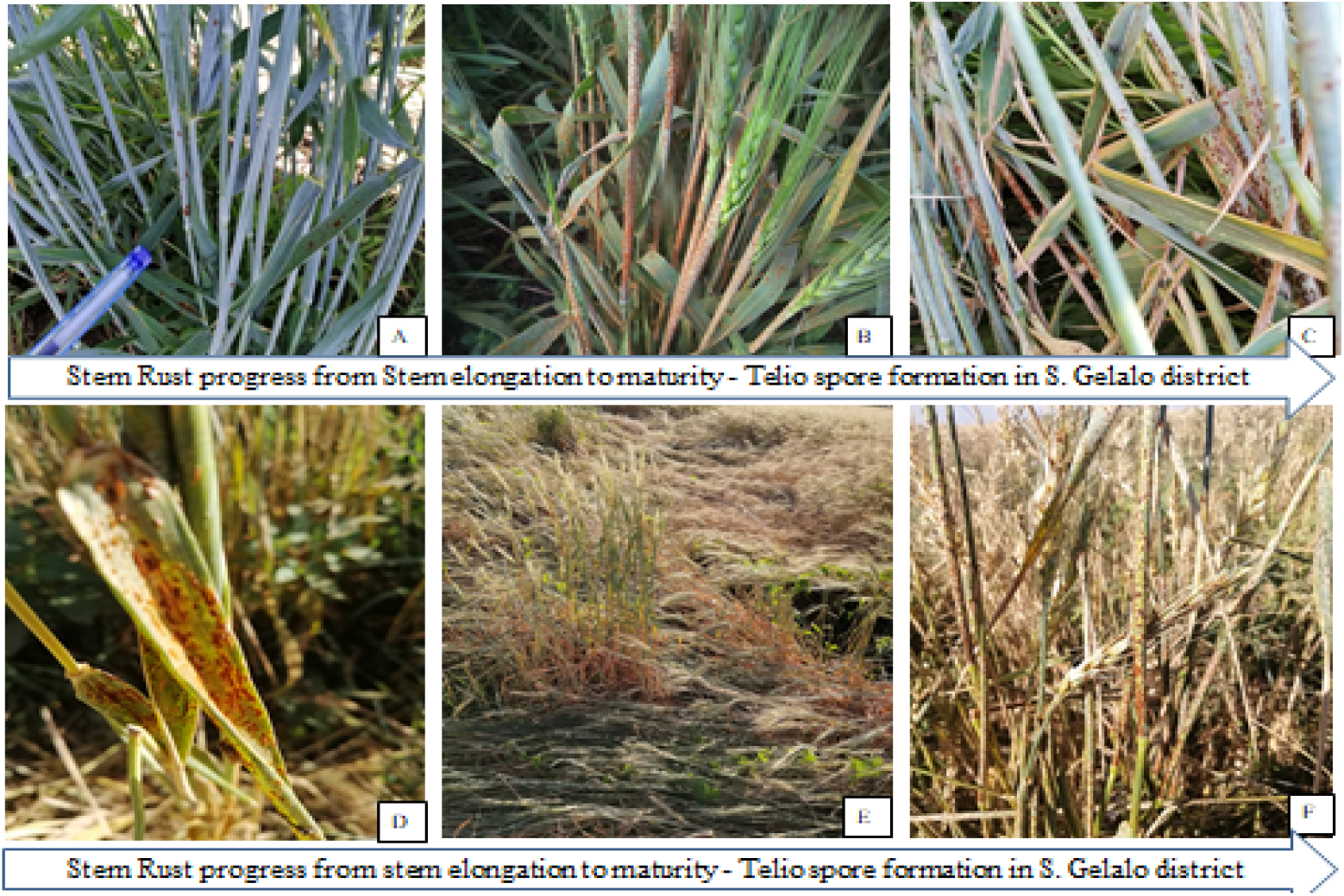
Stem rust virulence progress in a different stage of the plant: (A = stem elongation, B= Boating stage, C= milky stage, D= dough stage, E= matured, F= Teliospor formed in straw) districts of S. Gellalo districts

### 2.2. Survey of Wheat Stem Rust Distribution in Awash River Basin Fields

The survey and surveillance assessment were done among the farmers’ fields at every 5-10 km intervals following the main road routes in wheat farms. Stem rust assessment was carried out along the double diagonal ‘pattern and samples were taken using 1m^2^ quadrate. In each field wheat plants within the quadrate were counted and recorded as infected and healthy, and the intensity of stem rust was calculated. The prevalence, incidence, and severity of the diseases were calculated with the formula given below.

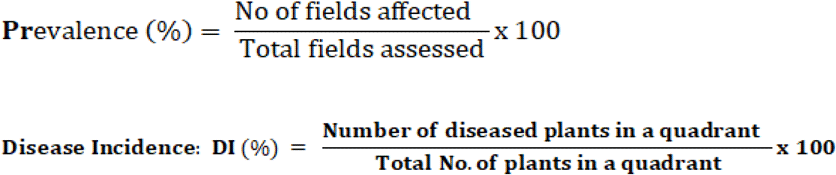

The disease severity under field condition was recorded followed by the modified Cobb’s scale as developed by Peterson *et al*. (1948).

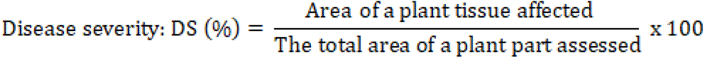

### 2.3. Seasonality and Spatial Epidemics Dynamics

The data on the prevalence, incidence, severity, and that of physiological races virulence spectrum were compared to each other to see the variations in season, and space in the Awash River basin irrigated wheat.

### 2.4. Physiological Race Identification of *P. graminis* f. sp. *Tritici*

#### 2.4.1. Collection of wheat stem rust samples

Samples of infected stems one sample per field were collected from wheat fields and trial plots in Awash River basin Oromia and Afar regional state of Ethiopia, to the period 2018/19-2019/20. Stems infected with wheat stem rusts were cut into small pieces of 5-10 cm using scissors and put in paper bags after the leaf sheath was separated from the stem to keep the stem and/or leaf sheath dry. During sample collection, we used absolute alcohol to sterilize the scissors to avoid cross-contamination among isolates. Samples collected in the paper bags were tagged with the name of the zone, district, longitude, latitude, variety, sample collection date, and transported to Ambo Agricultural Research Centers (AARC) for race analysis. The air-dried samples within the paper bags were kept in a refrigerator at 4°C until the survey and sampling in all districts are completed. Five seedlings of this variety for each sample were raised in suitable 8cm diameter clay pots that were filled with a mixture of steam-sterilized soil, sand, and manure in the ratio of 2:1:1, respectively. Spores were collected in the capsule from stem rust infected samples using a collector. The collected Urediniospores from each field were suspended in lightweight mineral oil, Soltrol 170, and inoculated using atomized inoculator on 7-day-old seedlings of variety McNair, which does not carry known stem rust resistance genes to get enough amount of spore to inoculate the stem rust differentials (Roelfs *et al*., 1992). When the pustules developed at two weeks spores from each pustule were collected using a power-operated vacuum aspirator and stored separately in gelatin capsules. A suspension, prepared by mixing urediospores with Soltrol 170, was inoculated on seven-day-old seedlings of the susceptible variety McNair for multiplication purpose for each of the single pustules on separate pots following the procedures mentioned earlier. The urediniospores descending from one pustule made up a single pustule isolate. One isolate was developed from each wheat field and used for the final race analysis (Roelfs *et al*., 1992).

Greenhouse inoculations were done using the methods and procedures developed by Stakman *et al*. (1962). The mono-pustule was inoculated, and the plants were then moistened with fine droplets of distilled water produced with an atomizer and placed in dew chamber for 18h dark at 18 to 22°^C^ followed by exposure to light for 3 to 4h to provide the condition for infection and seedlings were allowed to dry for about 2h. Then, the seedlings were transferred from the dew chamber to the growth room in the greenhouse where conditions were regulated at 12h photoperiod; at a temperature of 18 to 25°C and relative humidity of 60 to 70%.

Inoculated seedlings were moistened with fine droplets of distilled water produced with an atomizer and placed in a dew chamber in darkness for 18 h at 18 to 22°C and 98 to 100% relative humidity. Upon removal from the chamber, plants were exposed to 4 hours of fluorescent light to provide a condition for infection and allowed to dry their dew for about 2h. Inoculated plants were then transferred to greenhouse benches where conditions were regulated at 12h photoperiod, at the temperature of 18-25°^C^, and relative humidity (RH) of 60 to 70% (Stubbs *et al*., 1986). After seven to ten days of inoculation the flecks or chlorosis were visible leaves with a single fleck that produce a single pustule (single uredinal isolate) were selected from the base of the leaves and the remaining seedlings within the pots were removed by scissors. Only leaves with single pustules from each location were separately covered with Cellophane bags and tied up at the base with a rubber band to avoid cross-contamination (Fetch and Dunsmore, 2004).

#### 2.4.2. Inoculation of differential lines and race determination

Five seeds for each of the twenty wheat stem rust differentials with known stem rust resistance genes (Sr5, Sr6, Sr7b, Sr8a, Sr9a, Sr9b, Sr9d, Sr9e, Sr9g, Sr10, Sr11, Sr17, Sr21, Sr24, Sr30, Sr31, Sr36, Sr38, SrTmp, SrMcN, and a susceptible variety McNair) were grown in 10cm diameter pots. The susceptible variety ‘McNair’ (without Sr gene) was used to ascertain the viability of spores inoculated to the differential hosts. Each rust isolate derived from a single pustule was suspended in Soltrol 130. The suspension was adjusted to 3-5 mg urediospores with 1ml of mineral oil (solTrol-130) and inoculated onto seedlings of the differentials following the procedure described above. After inoculation, plants were moistened with fine droplets of distilled water produced with an atomizer and placed in an incubation chamber for 18 hours dark period at 18-22°C and 3-4 hours of light. Upon removal from the dew chamber, plants were placed in separate glass compartments in a greenhouse to avoid contamination and produce infection. Greenhouse temperature was maintained between 18°C and 25°C. Natural daylight was supplemented for additional 4 hours/day with 120 μ E.M^−2^ S^−1^ photosynthetically active radiations emitted by cool white fluorescent tubes arranged directly above plants to favor the pathogen produce infection.

Stem rust infection types (ITs) on the differential lines were scored 14 days after inoculation using a 0 to 4 scale (Stakman *et al*., 1962). Five letters race code nomenclature system was used according to Roelfs and Marten, (1988); Jin *et al*., (2008); and Hei *et al*., (2018). In this system the differential lines are grouped into five sub-sets, as shown in Table 1, in the following order:

i. Sr5, Sr2l, Sr9e, Sr7b
ii. Sr11, Sr6, Sr8a, Sr9g,
iii. Sr36, Sr9b, Sr30, Sr17.
iv. Sr9a, Sr9d. Sr10, SrTmp,
v. Sr24, Sr31, Sr38, SrMcN (Table 1).

**Table 1.** List of twenty wheat stem rust differential hosts with their corresponding Sr genes & origin/pedigree

| Differential hosts | Sr genes | Origin/Pedigree |
| --- | --- | --- |
| **LcSr24Ag** | 24 | Little Club/Agent (Cl 13523) |
| **W2691SrTt-1** | 36 | Cl12632 T. timopheevii |
| **ISr7b-Ra** | 7b | Hope/Chinese Spring |
| **ISr8a-Ra** | 8a | Rieti/Wilhelmina//Akagomughi |
| **CnSSrTmp** | Tmp | Triumph 64(Cl 13679)/ Chinese Spring |
| **Sr31(Benno)/ 6\*LMPG** | 31 | Kavkaz |
| **CnS-T-mono-deriv** | 21 | Einkorn Cl 2433 |
| **Trident** | 38 | Spear*4/VPM (Pl519303) |
| **ISr9a-Ra** | 9a | Red Egyptian/Chinese Spring |
| **ISr9d-Ra** | 9d | Hope/Chinese Spring |
| **Combination VII** | 17 | Esp 518/9 |
| **ISr5-Ra** | 5 | Thatcher/Chinese Spring |
| **ISr6-Ra** | 6 | Red Egyptian/Chinese Spring |
| **W2691Sr9b** | 9b | Kenya 117A |
| **Vernsteine** | 9e | Little Club//3*Gabo/2* |
| **W2691Sr10** | 10 | Marquis*4/Egypt A95/2/2*W2691 |
| **BtSr30Wst** | 30 | Festival/Uruguay C10837 |
| **CnsSr9g** | 9g | Selection from Kubanka(Cl1516) |
| **ISr11-Ra** | 11 | Kenya C6402/Pusa4/Dundee |
| **McNair 701** | McN | Cl 15288 |

Source; Ambo plant protection research center, 2009

An isolate that produces a low infection type on the four lines in a set is assigned with the letter ‘B’, In comparison, a high infection type on the four lines is assigned with the letter ‘T’. Hence, if an isolate produces a low infection type (resistant reaction) on the 20 differential lines, the race could be designated with a five-letter race code ‘BBBBB’ while an isolate producing a high infection type could be designated with a five letters ‘TTTTT’ and race analysis was by using the descriptive statics.

**Table 2.**
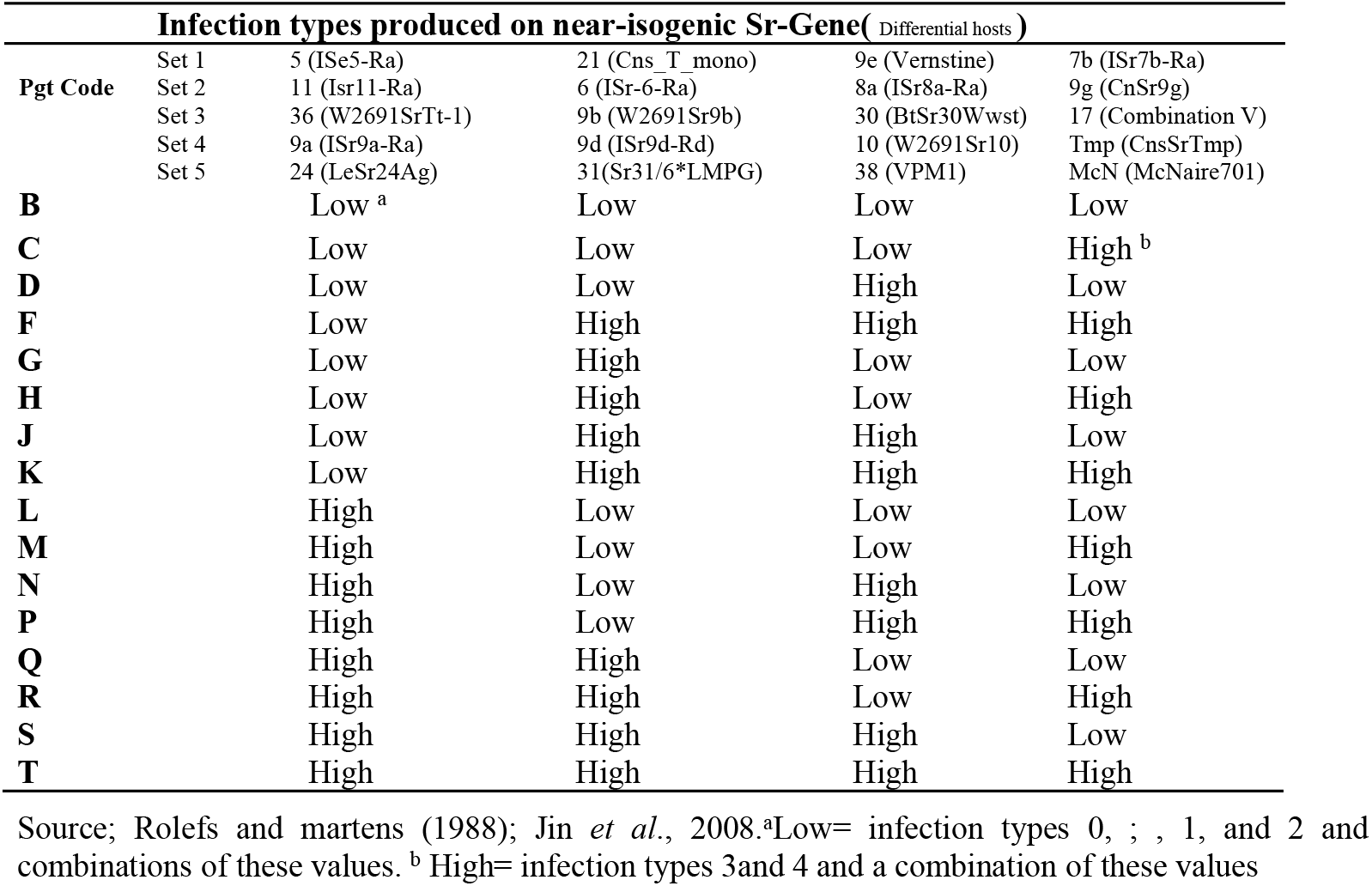
Wheat stem rust differentials and Nomenclature of Pgt based on 20 ** wheat hosts

##### Dead Stem Rust DNA Samples

Stem rust DNA analysis was done from samples collected in 2019. A total of 9 samples were collected from widely dispersed and representative locations in irrigated wheat-growing areas. The isolate was collected by cutting immediately above and below a single pustule using sharp scissors. The sheath tissue was cut at the side and the stem was removed. The samples were kept in 80% alcohol for seven days. After pouring the alcohol the samples were left open to dry for two days. Then the samples were sealed tubes shipped to CDL Minnesota for DNA analysis.

### 2.5. Data Analysis

Survey data (prevalence, incidence, and severity) were analyzed by using descriptive statistical analysis (means) over districts, varieties, and altitude range, and crop growth stages. Similarly, race analysis was analyzed using those descriptive statistics (Gomez and Gomez, 1984).

## 3. RESULTS

### 3.1. Distribution of Stem Rust in Awash River Basin

During 2014/15, 2015/16, and 2017/18 stem rust was intercepted in single farm spots, and in 2016 there was no detection of stem rust epidemics at all studied locations. This means the diseases stem rust was not economically important in those seasons on wheat variety ‘Gambo’ This could be due to the production areas were new for the stem rust host plant wheat and the introduction of spores was not established or maybe those production seasons were not conducive for rust outbreak. In 2018/19, a total of 33 stem rust live samples were taken, and only (12.12%) samples were viable, while in 2019/20 a total of 46 stem rust live samples were taken and (71.74%) samples were viable. From a total of 79 stem rust samples taken for the two seasons, (46.84% of) samples were viable which is low overall viability could be associated with loss of sample viability with storage environmental condition. The low viability of samples could be attributed to the late arrival of the collected samples or sample miss handling. The spatial viability percentage to collected samples in 2019/20, in Oromia regional state, were 5(40%), 6(100%), and 17(70.59%) in Sire, Jeju, and Fentale districts respectively; While on Afar regional state (33.33%) 6, (100%) 8 and (25%) 4 in Afambo, Amibara and Dubti districts respectively were viable.

In 2014/15, the number of fields assessed was three in Amibara and seven in Fentale of which 1 farm was affected with stem rust at physiological maturity. Even though the race analysis was not done the first record of the disease was in Fentale district Gedara Kebele in 2014/15 production year with prevalence (14.3), Incidence (1.4), and Severity (0.7) in percentage. In 2015/16 the stem rust expansion increased to a prevalence of 20%, 1% incidence, and 1.5% severity in the Fentale district. During the same year, stem rust infestation coincided with late dough stage, such that the diseases were visible in ditches of undulated sites.

In 2016/17, stem rust pressure was not experienced in Awash River basin irrigated wheat valley. This could be attributed to sunny weather 15 December -15 January. In 2017/18 the stem rust started to invade the wheat production in Amibara with incidence (1.33%), severity (0.5%) although there was no disease detected in Fentale.

In 2018/19 the diseases escalated in terms of prevalence (71.7%), incidence (47.45%), and severity (29.27%) in Awash River basin. The highest incidence of stem rust was noted in Gellalo with (83.33%), the second-largest incidence was in Fentale (65%), and the third incidence was in Amibara (60%). In Sire and Jeju districts of the Oromia region the incidence of Sr was 30% and 50% respectively, whereas the lowest incidence was in Dubti (;) during physiological maturity. The disease severity showed a similar trend as the incidence. The highest severity in Gellalo with range mean values 60-80%, (70%) respectively. This was followed by Fentale and Amibara districts with a range of 0-80% for both districts and a mean value of (40%) and (35.63%) respectively. In Sire and Jeju districts, the severity was (16.67%) and (28.5%) respectively. The rust-free district was Gewane due to poor management performance of the wheat was also bad, due to management after booting they stopped irrigation water completely. Even though the late planting date increases the outbreak of stem rust diseases. Stem rust is polycyclic by its nature in those types of air-borne diseases the spore pressure in the air was critical to the nature of disease virulence. Most of the varieties released in Werer ARC including Fentale two were susceptible to stem rust. Fentale two ranges with 100% incidence and 80% severity in Werer Seed multiplication leased farm. The losses detected with this disease was also affected the seed multiplication unit of the center and caused high yield loss. In 2018/19 the disease invades most wheat production districts in the basin and incurs economic losses in terms of agricultural input cost for fungicide purchases, application cost, and yield losses. The disease pressure was severing in farms near to Awash River with high morning dew pressure and low on farms which are far from riverbanks.

The pressure of the disease was more severe in the farms near Awash River with high morning dew pressure and low on farms that are far from riverbanks.

**Table 3.** Prevalence, incidence, and severity of stem rust in two districts of Awash River Basin in 2014/15

| Region | District | Altitude range (m) | NFI | P (%) | I (%) |  | S (%) |  |
| --- | --- | --- | --- | --- | --- | --- | --- | --- |
|  |  |  |  |  | Range | Mean | Range | Mean |
| **Afar** | Amibara | 735- 742 | 3 | 0 | - | - | - | - |
| **Oromia** | Fentale | 1030-1045 | 7 | 14.3 | 0-10 | 1.4 | 0-5 | 0.7 |
| **Total** |  | 735-1045 | 10 | 10 | 0-10 | 1.0 | 0-5 | 0.5 |

NFI=No. Diseased Field Inspected; P=Prevalence I= incidence; S= Severity

In 2014/15 prevalence and intensity of stem rust in small-scale producers’ fields were negligible in the Awash River basin; the first intercept in irrigated wheat production in Ethiopia was in the Fentale district for the year. The most probable reason for low disease intensity was a low source of inoculum or the pathogen in the new production area.

**Table 4.**
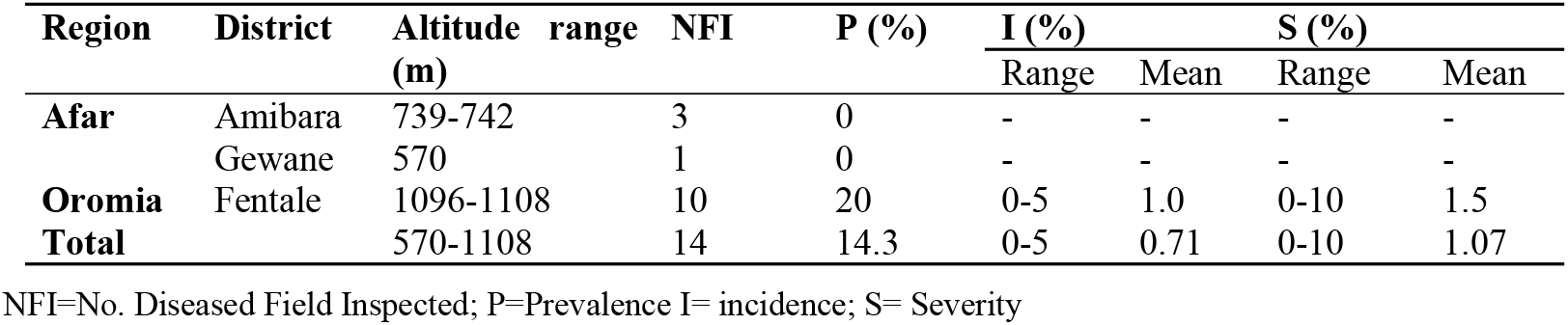
Prevalence, incidence, and severity of stem rust in three districts of Awash River Basin 2015/16

In 2015/16, Wheat production was at small scale farmers of Amibara, Gewane, and Fentale districts the prevalence and intensity of stem rust in small scale producers in awash basin fields were still low; with an increment of the 10 fields prevalence in Fentale districts from 14.3% to 20.0% the intensity was still negligible.

**Table 5.** Prevalence, incidence, and severity of stem rust in two districts of Awash River Basin 2016/17

| Region | District | Altitude range (m) | NFI | Prevalence (%) | Incidence (%) |  | Severity (%) |  |
| --- | --- | --- | --- | --- | --- | --- | --- | --- |
|  |  |  |  |  | Range | Mean | Range | Mean |
| **Afar** | Amibara | 741-742 | 2 | 0 | - | - | - | - |
| **Oromia** | Fentale | 1090-1104 | 8 | 0 | - | - | - | - |
| **Total** |  | 741-1104 | 10 | 0 | - | - | - | - |

NFI=No. Diseased Field Inspected; P=Prevalence I= incidence; S= Severity

**Table 6.**
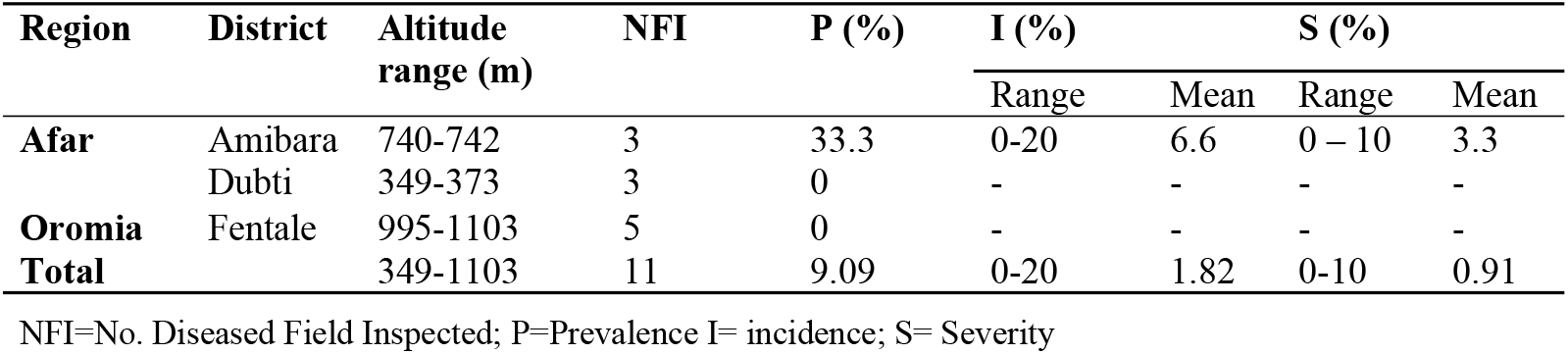
Prevalence, incidence, and severity of stem rust in three districts of Awash River Basin 2017/18

In 2017/18 there was no stem rust disease prevalence in irrigated lowlands.

**Table 7.**
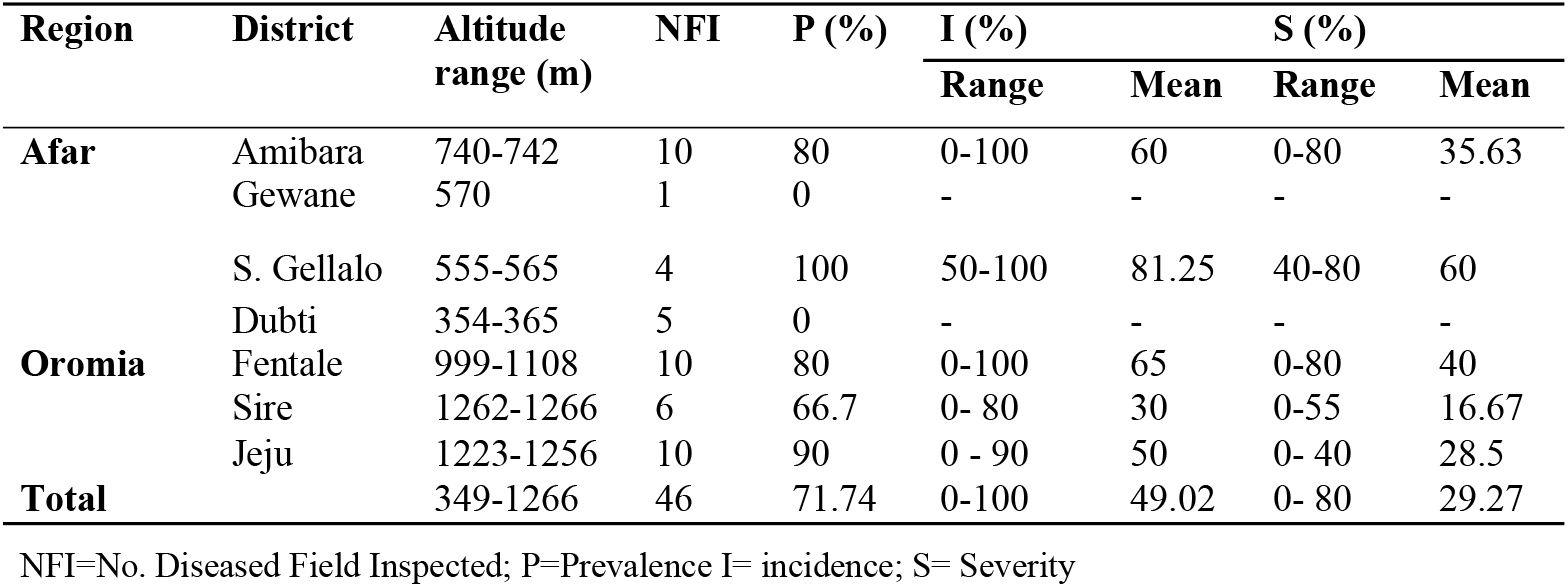
Prevalence incidence and severity of Stem rust in seven districts of Awash River Basin (2018/19)

In 2018/19 wheat production was expanded to large-scale producers and sugar corporation farms. The Pgt prevalence reached 80% in Amibara 100% in Gellalo districts of middle awash and there was no Pgt infection in lower awash sugar corporation farms at Dubti. While the prevalence of the diseases was 80% in Fentale, 90% in Jeju, and 66.7% in Sire districts of Oromia.

**Table 8.**
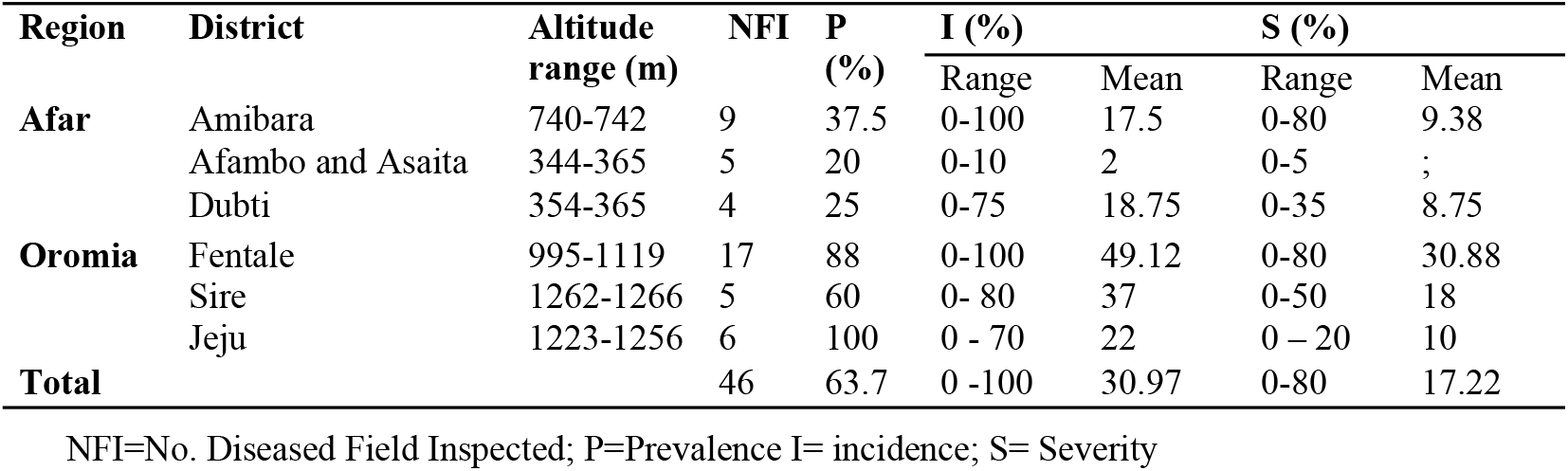
Prevalence incidence and severity of Stem rust in seven districts of Awash River Basin (2019/20)

### 3.2. Stem Rust Disease Dynamics in Awash River Basin

The Pgt dynamics in the low land areas its disease measuring parameters and infection increased from production year to year.

The Pgt disease prevalence as shown above in figure (3A) was increasing from time to time with the expansion of wheat production and reached 80% prevalent in the 2018/19 production year on upper awash and middle awash areas. In the 2019/20 production year, the prevalence of the wheat stem rust was more prevalent in upper awash 85.6% in upper awash; and the prevalence of the rust was about 30% in middle awash about 20% prevalence was observed in lower awash farmer and sugar corporation seed multiplication fields.

**Figure (3).**
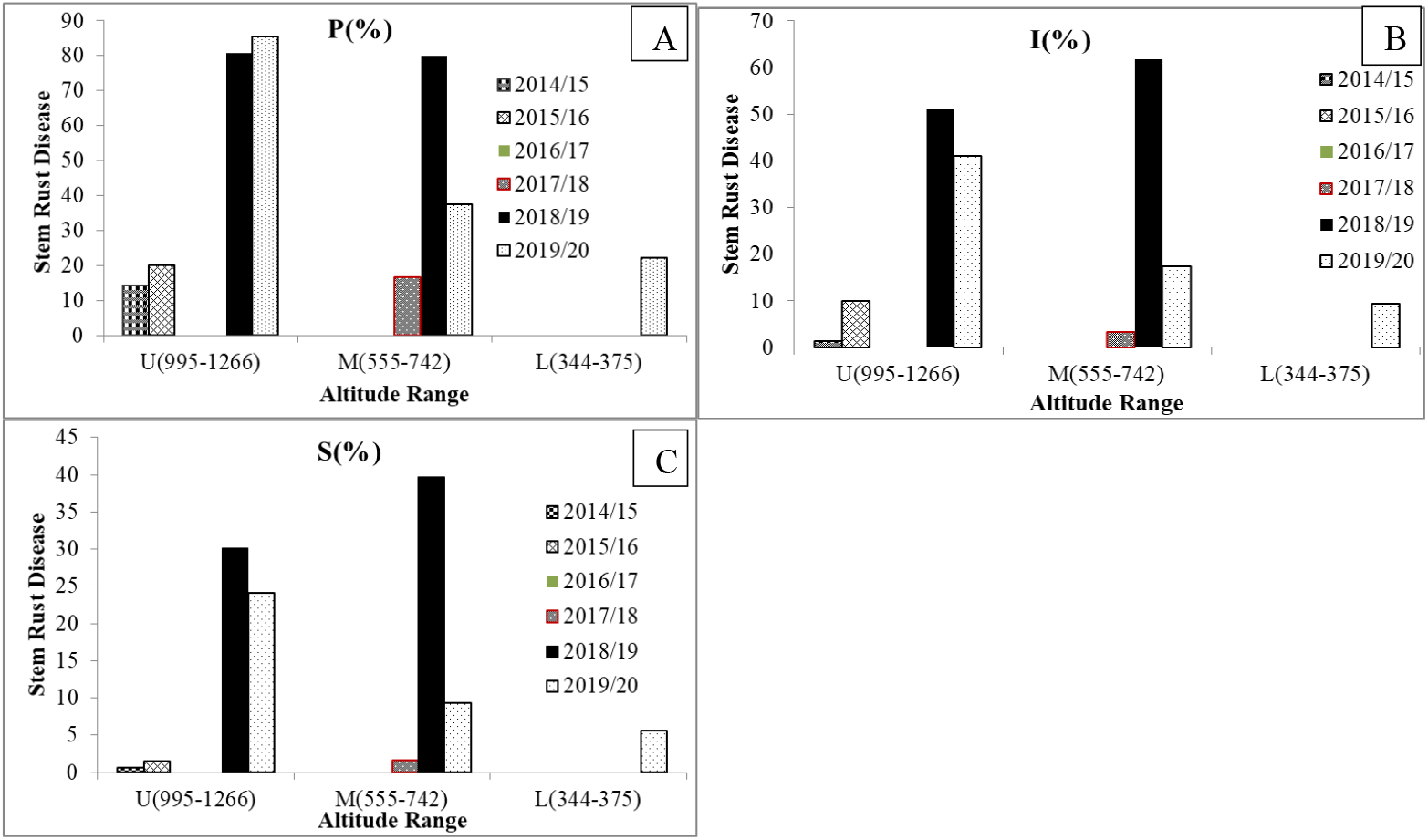
A (P%) = Prevalence, B (I%) = Incidence, and C (S%) = Severity percentages of stem rust disease in with three altitude range of Awash Basin areas considered of; U=Upper Awash, M= Middle Awash, and L= Lower Awash.

*Pgt* disease intensity was highest in Middle Awash than Upper Awash and Lower Awash which is healthy for 2018/19. The incidence of *Pgt* as depicted above in figure (3B) was increasing from year to year. The result indicated that *Pgt* disease intensity in the Awash River basin was majorly influenced by weather factors. In 2018/19 the weather condition of the area was cloudy which is conducive for stem rust. Pgt incidence in 2018/19 was higher in Middle Awash than on Upper Awash. Generally speaking, the disease pressures were higher on Upper and Middle Awash than Lower Awash. That could be possibly related to the environmental condition in Lower Awash i.e. hot sunny with high evapotranspiration which removes the dews before germination of spores to the occurrences of disease infection.

In 2018/19 of the 46 fields, 56.5% of the fields surveyed were found at an altitude range of (995-1266) m.a.s.l. also known as the upper awash, 32.61%, of the field surveyed, were at an altitude range of (555-742) m.a.s.l. also known as middle awash and the rust 10.87% was at altitude range (344-375) m.a.s.l. also known as lower Awash in Awash basin classification. The data showed that the number of stem rust-infected fields decreased as the altitude decreased. The incidence also decreased from 51.15% at upper awash to 61.66% at middle awash and 0% at lower awash. The same results were recorded regarding the disease severity; 32.76% at upper awash 39.75% at middle awash and 0% at lower awash fields. In 2019/20 of the 46 fields, 60.86% of the fields surveyed were found at an altitude range of (995-1266) m.a.s.l., in upper awash, 19.57%, of the field surveyed were at an altitude range of (555 - 742) m.a.s.l., in middle awash 19.57% of fields were at altitude range (344-375) m.a.s.l. and in lower Awash in Awash basin classification.

The data showed that the number of stem rust-infected fields decreased as the altitude decreased. The incidence also decreased from 41.14% at upper awash to 17.5% at middle awash and 8.33% at lower awash. The same result was recorded regarding the disease severity 24.11% at upper awash 9.38% at middle awash and 3.89% which is a trace in lower awash fields. In rain-fed agroecology Ayele *et al*., (2008) reported that the highest level of stem rust infection was recorded in the altitude range of 1600 – 2500. High humidity along with high temperatures during the growing season in the low altitude areas favors the growth of the pathogen (Roelfs *et al*., 1992). In cool-season irrigated lowlands wheat production stem rust of wheat was very important in altitude range above 555 m.a.s.l middle and upper awash areas. Our result strengthens the findings of the above authors and the environmental requirement of rusts determined by the microclimate. Generally, the stem rust disease intensity increases in the middle and upper awash areas of Awash River basin, and the initial infection was started in Lower Awash in 2019/20 as depicted in the graph below (Figure 4).

**Figure 4.**
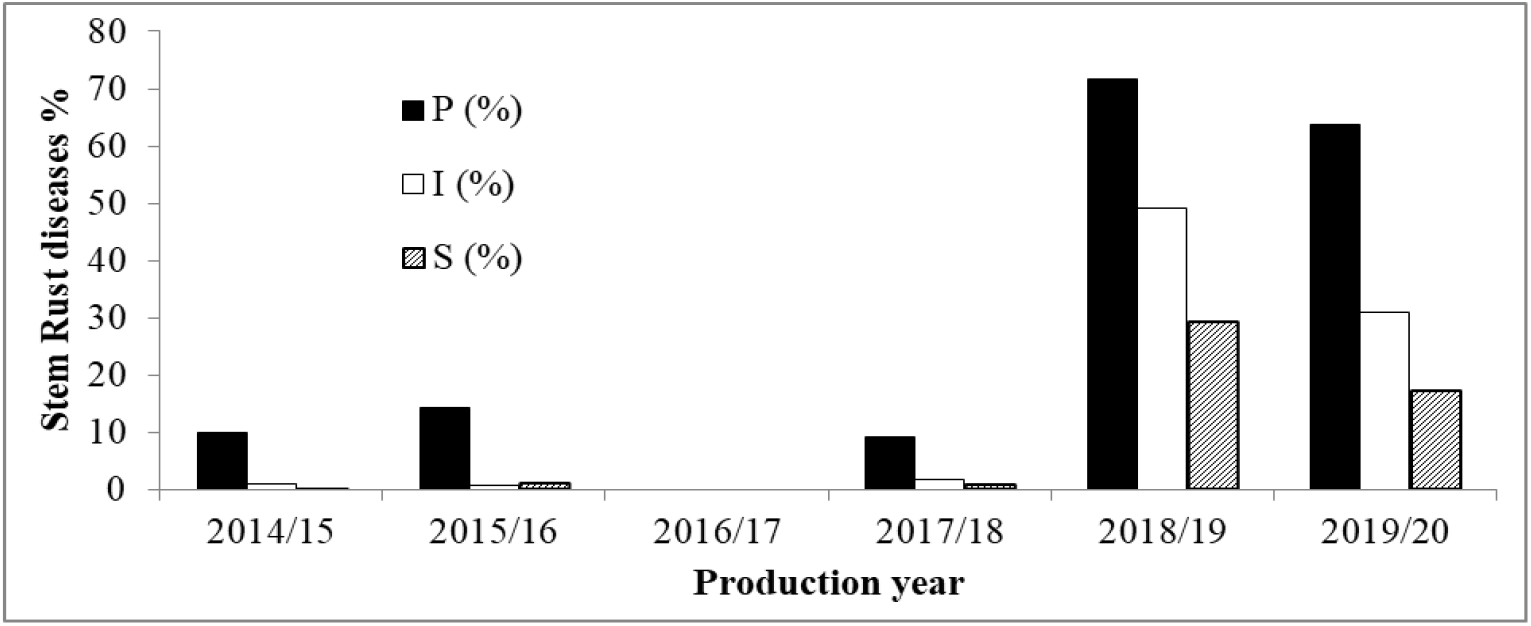
Prevalence, incidence, and severity of stem rust dynamics year to year of the areas considered

Stem rust diseases in the Awash River basin were insignificant and quite low like its production up to 2017/18. The disease infection was quite increasing in the 2018/19 and 2019/20 production years as the progress of new area production coverage is expanding.

### 3.1. Virulence and Physiological Races of Stem Rust Pathogen

Distribution and frequency of Pgt. races in Awash River basin for 2014/15 - 2017/18 off seasons in irrigated wheat cropping season there was no race analysis done before.

In the year 2016, there was no detection of stem rust in the Awash River basin which could be a result of poor crop management with unsuitable weather conditions i.e., sunny in the area. In 2018/19 a total of 33 stem rust samples were taken to the Ambo Agricultural research stem rust laboratory. Only Four samples as different races were viable due to unknown reasons (12.12%) viability of samples were quite low due to unknown reasons related to the late arrival of the collected samples.

Surveys in 2019/20 a total of 46 stem rust samples were collected and they were taken AARC for physiological race analysis. During this cropping season, a total of 28 stem rust samples in Oromia (Sire, Jeju, and Fentale) districts and 19 samples were collected from Afar regional state (Afambo, Dubti, and Amibara) districts.

In 2019/20 a total of 46 stem rust samples were taken and 33 (71.74%) samples were viable the rest 13 (28.26%) were not viable due to unknown reasons which are obvious in many cases with mistakes during sample handlings; Among the collected samples in Sire 2(5), Jeju 6(6) and Fentale 12(17) districts respectively; While in Afambo and Asaita 2(6), Dubti 1(4) and Amibara 8(8) were viable. Urediniospores collected from each field were suspended in lightweight mineral oil, Soltrol 170, and inoculated using an atomized inoculator on 7-day-old seedlings of variety McNair, which does not carry known stem rust resistance genes (Roelfs *et al*., 1992).

**Table 9.**
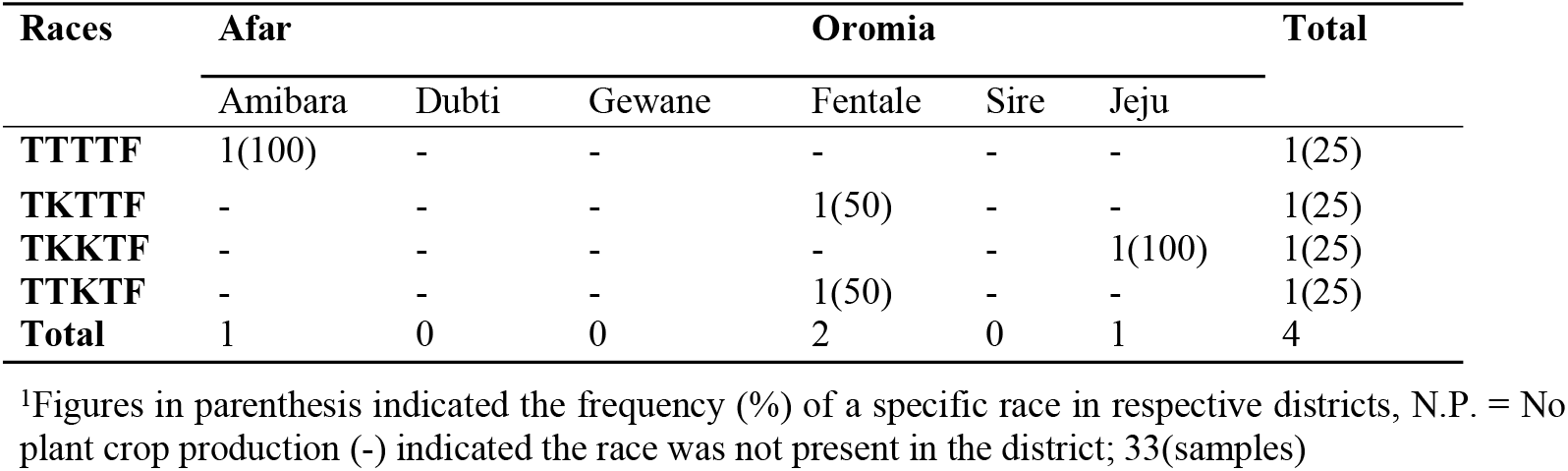
Distribution and frequency of P.gt. races in Awash River Basin Oromia and Afar regions of Ethiopia for 2018/19 for the irrigated wheat cropping season

The Pgt race analysis result indicates that the

**Table 10.** Distribution and frequency of P.gt. races in Awash River basin irrigated wheat in 2014 and 2019/20 main cropping season

| Races | Afar |  |  | Oromia |  |  | Total |
| --- | --- | --- | --- | --- | --- | --- | --- |
|  | Gebiresu | Zone #1 |  | East Shewa | East Arsi |  |  |
|  | Amibara | Dubti | Afambo | Fentale | Sire | Jeju |  |
| TTTTF | 5(62.5) | - | - | 5 (41.7) | 2(66.6) | 3(42.9) | 15(45.5) |
| TKTTF | 2(25) | - | - | 3 (25) | - | - | 5(15.2) |
| TKKTF | 1(12.5) | 1(100) | 2(100) | 2 (16.7) | - | 2(28.6) | 8(24.2) |
| TTKTF | - | - | - | 1 (8.3) | 1(33.3) | 2(28.6) | 4(12.1) |
| TKPTF | - | - | - | 1 (8.3) | - | - | 1(3.0) |
| Total | 8 | 1 | 2 | 12 | 3 | 7 | 33 |

<sup>1</sup>Figures in parenthesis indicated the frequency (%) of a specific race in respective districts, N.P. = No plant crop production (-) indicated the race was not present in the district; 33 (46) (samples)

**Table 11.**
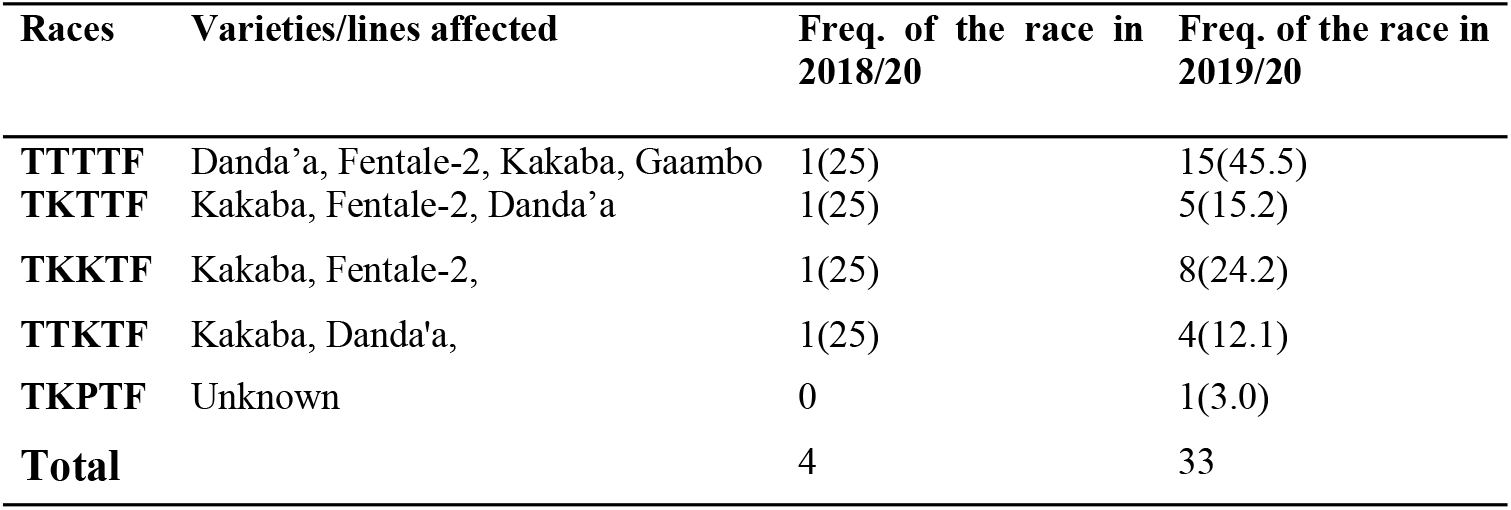
Irrigated Wheat varieties were affected by the Pgt races identified in 2018/19 and 2019/20 cool seasons and their frequency from the total viable Pgt population in the areas.

| Races | Varieties/lines affected | Freq. of the race in 2018/20 | Freq. of the race in 2019/20 |
| --- | --- | --- | --- |
| TTTTF | Danda'a, Fentale-2, Kakaba, Gaambo | 1(25) | 15(45.5) |
| TKTTF | Kakaba, Fentale-2, Danda'a | 1(25) | 5(15.2) |
| TKKTF | Kakaba, Fentale-2, | 1(25) | 8(24.2) |
| TTKTF | Kakaba, Danda'a, | 1(25) | 4(12.1) |
| TKPTF | Unknown | 0 | 1(3.0) |
| **Total** |  | 4 | 33 |

The figure in the parenthesis indicated the frequency (%) of a specific race in the season.

Below is the result from CDL Minnesota. Also, find attached the raw data from the survey along with the result of the genotyping. According to the genotyping from the Minnesota CDL lab, all 9 samples were genotyped as clade III-B (Co-A04) which is associated with *Pg*t race TTRTF. These results are consistent with race phenotyping work showing that the Pgt races TTRTF (III-B) and TKFTF/TKKTF (IV-F) have become the prominent races in Ethiopia.

**Table 12.** a virulence or virulence spectra of the Pgt races identified in 2018/19 and 2019/20 cool seasons irrigated wheat areas

| Races | Virulence | Avirulence |
| --- | --- | --- |
| TTTTF | 5, 21, 9e, 7b, 11, 6, 8a, 9g, 36, 9b, 30, 17, 9a, 9d, 10, Tmp, 38, McN | 24, 31, |
| TKTTF | 5, 21, 9e, 7b, 6, 8a, 9g, 36, 9b, 30, 17, 9a, 9d, 10, Tmp, 38, McN | 11, 24, 31 |
| TKKTF | 5, 21, 7b, 6, 8a, 9g, 9b, 30, 17, 9a, 9d, 10, Tmp, 9e, 38, McN | 11, 36, 24, 31 |
| TTKTF | 5, 21, 9e, 7b, 11, 6, 8a, 9g, 9b, 30, 17, 9a, 9d, 10, Tmp, 38, McN | 36, 24, 31 |
| TKPTF | 5, 21, 9e, 7b, 6, 8a, 9g, 36, 30, 17, 9a, 9d, 10, Tmp, 38, McN | 9b, 11, 24, 31 |

#### Virulence to Sr genes

About 100% of the races identified showed virulence above the range of 80% of the Sr genes. The races TKKTF and TKPTF showed virulence to 80% of the Sr genes. The races TKTTF and TTKTF showed 85% of Virulence. The race TTTTF showed virulence to 90% of Sr genes. The race TTTTF was virulent to all differential lines except Sr24 and Sr31. Races TKTTF and TTKTF differed by only one gene, which was virulent to all differential lines except Sr11, Sr24, Sr36, and Sr31, Sr24, Sr36, respectively. The virulence pattern observed in this study conforms to the results reported by Naod *et al*. (2005); Belayenh and Emebet (2005), and Admassu *et al*. (2009).

This study showed that the stem race races in the irrigated wheat production areas of the Awash River basin were highly virulent to most of Sr genes. The only effective genes able to provide a good harvest in the existing races are Sr 24 and Sr 31. On the other hand, more than 80% of the Sr genes were defeated by 100% of the isolates. The resistance gene McNair (Sr McN) was ineffective to all isolates tested (Table 4). During 2018/19 a total of 33 isolates were collected in cool seasons. Out of this only 4 were viable and analyzed on to the 20 stem rust differential lines in 2018/19. From the viable samples, four isolates were studied in the season and four different races namely TTTTF, TKTTF which is also known as Digelu race, TKKTF, and TTKTF were identified. The result was in agreement with the findings of Hei *et al*. (2018) who noted that, about 95% of the stem rust population was TTTTF and which was identified from durum wheat, bread wheat, and Barley varieties during 2014-2015.

## 4. Discussion

Irrigated Wheat crop production is grown in cool crop growing season with improved crop varieties in Awash River basin of Ethiopia. In the basin, wheat production has been increasing to substantial levels to support wheat self-sufficiency. On the other hand, stem rust disease prevalence, incidence, and severity have been increasing and the disease may reach a level that threatens wheat production and productivity. Therefore, it is essential to take appropriate action to avert a disaster from occurring.

The wheat production trend increases at an alarming rate with the intention of the government of Ethiopia to substitute the imports and stabilize the wheat self-sufficiency. Thus, a plan could become possible by deploying best-bet wheat production technologies (MOA, 2020). The production expansion to potential irrigated areas of Ethiopia was in fast progress and there is a huge plan to cover the basins and used the water resources with the existing irrigable lands of the country including the Abay river also known as the Nile River in the northwest highlands and western Ethiopia foothill lowlands in the basin. With our little experience for the past six years in area expansion and cropping frequency, one of the potential problem or challenge which needs a strategic intervention were biotic stress majorly stem rust.

Stem rust is a major disease in irrigated wheat belts of Ethiopia and particularly in the Awash River basin and they can cause significant yield losses and quality in years with suitable conditions. The intensity of the disease changes from year to year and from place to place depending on the area’s macro and microclimatic conditions. Its prevalence incidence and severity were increasing from year to year and with increasing area production. The disease was more important in farms near to Awash River than at the distant farms from the river. The soil type and the nature of the farm also contribute to the disease. Fields surrounded by mountain ranges and clay loam soils have high pressure than the sandy loam soil types. The cool season wheat production in lowlands for the majority of wheat fields assessed in the Awash River basin ranges from 360 - 1264 m.a.s.l. The disease was most prominent in the altitude range (560 - 1264) m.a.s.l. and its pressure was the peak in the range (560-1064) m.a.s.l. namely Gewane (50%), Fentale (35%), and Amibara (35.63%) severity in 2018/19. This result also shows the high susceptibility of the good adaptable variety Kakaba to stem rust. So the production of highly adaptable good-yielding varieties must be supported with the use/application of fungicide chemicals.

The 37 samples analyzed in both seasons the TTTTF, TKTTF, TKKTF, TTKTF, and TKPTF were identified. This indicated the presence of broad races with a wider virulence spectrum within the Pgt population in the irrigated areas. This is in line with previous studies conducted by (Belayneh and Emebet, 2005; Admassu *et al*., 2009; Hailu *et al*., 2015; Hei *et al*., 2019). This area was a new production corridor of wheat there is an expectation to find new races while the results showed that the races are similar to the main season production areas with the tendency of new race formation or introduction. Continuous wheat production and the favorable microclimate in Ethiopia irrigated wheat production areas could be the reason for the rapid evolution and high virulence diversity of the pathogen.

The most abundant dominant virulent race was TTTTF. It had frequencies of 25 and 45.45% of stem rust races for 2018/19 and 2019/20 irrigated wheat production areas Afar and Oromia regions of Ethiopia. The race TTTTF had a wide virulence spectrum it has similar virulence formula to TKTTF. The second abundant and dominant virulent race was TKKTF. It had frequencies of 25% and 24.24% in the 2018/19 and 2019/20 cropping seasons. This race was the only dominant virulent race in lower awash; TKKTF could have a large adaptation range for environmental factors. The third abundant and virulent race was TKTTF (Digalu race). It had frequencies of 25.0 and 15.15% in 2018/19 and 2019/20 cropping seasons, respectively. Races TTTTF, TKKTF, TKTTF accounted for 79.9% of the stem rust population of irrigated wheat production areas in the study period. The remaining two races were the least abundant TTKTF 18.8% and TKPTF 1.5%. The Race TKPTF was detected at a single location at the Fentale district of East Shewa in 2019/20. This study revealed that TTTTF was the most dominant and virulent of the stem rust populations in Oromia irrigated wheat agro-ecologies. While the race TKKTF was the most dominant and virulent of the stem rust population in the Afar region. The races were widely distributed in the major irrigated wheat growing areas of Ethiopia in the seasons. This result was in agreement with most of the recent studies in the rain-fed areas wheat production reports. As it was reported by Lemma *et al*. (2015), the major race of wheat stem rust detected in the East Shewa Zone of central Ethiopia was TTTTF followed by TKKTF. It was also reported in Iran (Afshari *et al*., 2015) and Italy (FAO 2017). Moreover, Yehizebalem *et al*. (2020) also reported that TTTTF, TKTTF, and TKPTF were found to be the stem rust race populations that were affecting the wheat production of North West Ethiopia.

## 5. CONCLUSION

Crop production in Ethiopia was started to shift from the rain feed dominated to intensive irrigated agriculture. However irrigated agriculture, its merit in climate change needs full production package skills to boom wheat production and productivity. For the last one decade the irrigated wheat production. The highest stem rust incidence and severity were observed in Gewane and Fentale districts. Whereas the lowest incidence and severity were recorded in lower awash Dubti districts. Stem rust samples were taken at altitude ranges of (560 to 1108 m.a.s.l.). The severity of the disease was relatively high at farms nearest to Awash Riverbanks and high dew point areas. The incidence and severity of the disease were higher on variety Fentale two than the others in research seed multiplication plots. The bread wheat variety Kakaba was used for its high adaptability and yield potential on all farmers’ fields. The highest stem rust intensity was observed during the dough growth stage than at an earlier growth stage. The collected survey samples were analyzed on the twenty standard stem rust differentials to the variability of the pathogen using physiological race analysis resulted in four races. The highly virulent race TTTTF was among the important race identified in the area. The other important races were TKTTF (Digalu race) TKKTF, TTKTF. Differential host carrying Sr 24 was an effective gene that results in resistance to 100% of the area’s races. Even though the long-distance he a high rate of invasion with a fast dispersal rate nature spores movement of virulent races like TTKSK or ug99, and its variant gene virulent to resistant wheat genes are expected. Besides, new races could be expected in unexploited potential areas that could affect the resistant varieties for those existing races.

## 6. ACKNOWLEDGEMENTS

we are privileged to deliver our warm gratitude to DGGW Project for the financial support without them this experience would never pass the nightmare. Heartedly, I am grateful to DGGW Project for the 2018 budget year financial support, and Cambridge University 2019/20 production year for both field and laboratory work. I am greatly indebted to the EIAR for allowing me to conduct the survey and Werer Agricultural Research Center, for the logistic supports. I am happy to thank the drivers for their patience and willingness in data collection in the warm weather. I would like to thank, Ambo wheat stem rust team Tizazu T. and Teklu N. for generating and providing physiological race analysis data. Last but not least, I would like to deliver my gratitude to Mr. Direba Megeressa for helping us in plotting the study Map.

**Appendix Table 1.**
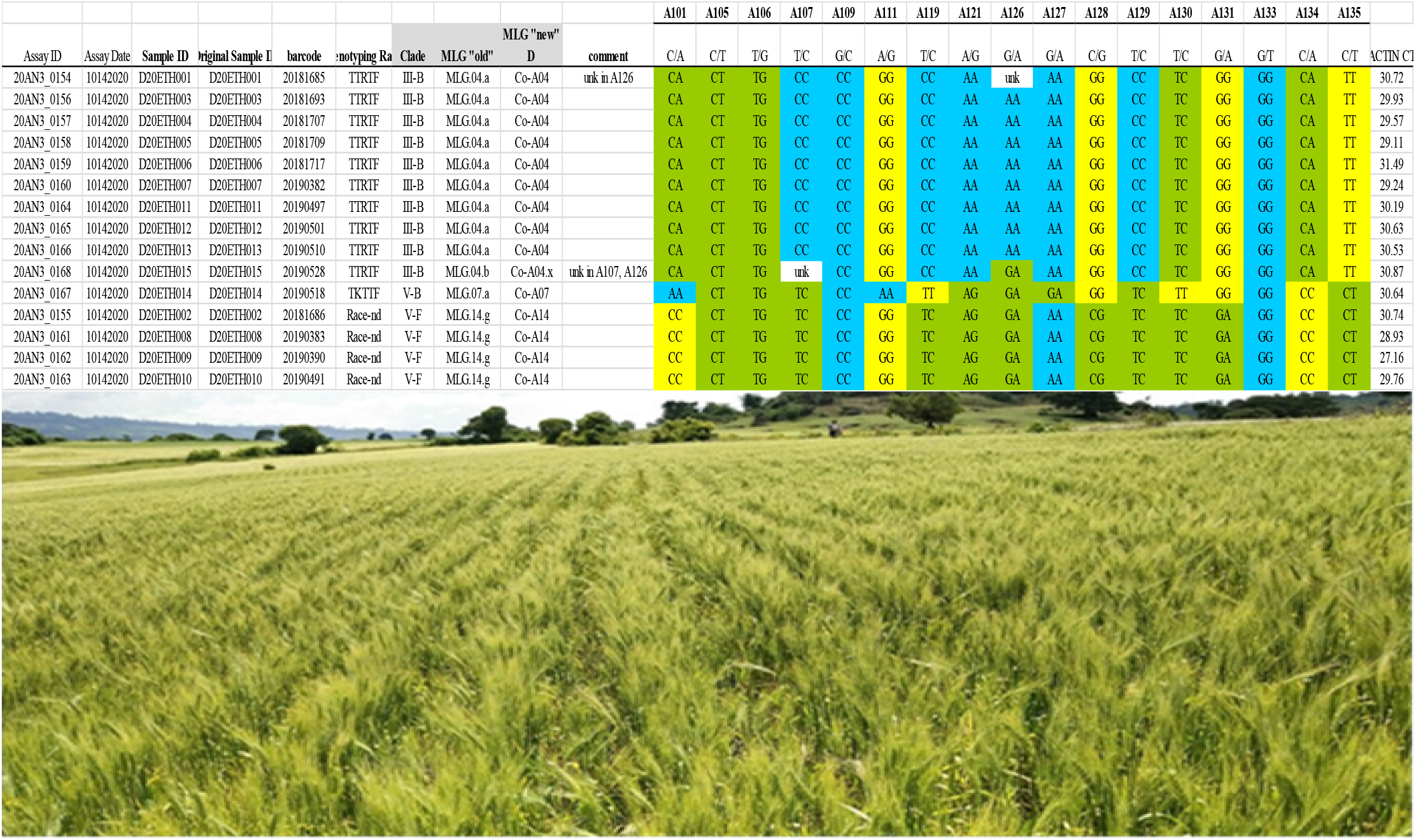
*Pg*t Genetic diversity in upper Awash River basin of Ethiopia; core SNP assay CDL results from Minnesota

## REFERENCE

Abebe, T., Woldeab G., and Dawit W., 2012. Distribution and physiologic races of wheat stem rust in Tigray, Ethiopia. Journal of Plant Pathology & Microbiology.

Admassu, B., and Fekadu E. 2005. Physiological races and virulence diversity of Puccinia graminis f. sp. tritici on wheat in Ethiopia. Phytopathologia Mediterranea, 44(3), pp.313–318.

Admassu, B., Lind V., Friedt W., and Ordon F., 2009. Virulence analysis of Puccinia graminis f. sp. tritici populations in Ethiopia with special consideration of Ug99.Plant Pathology, 58(2), pp.362–369.

Afshari, F., Aghaee M., Jalal Kamali M.R., Roohparvar R., Malihipour A., Khodarahmi M., EbrahimnejadSh A.R, Chaichi M., Dadrezaei S.T., Dalvand M. and Dehghan M.A., 2015. Surveillance and Pgt race analysis in Iran, 2014. Borlaug Global Rust Initiative, 123pp.

Amanuel, M., Gebre, D., and Debele, T., 2018. Performance of bread wheat genotypes under the different environments in lowland irrigated areas of Afar Region, Ethiopia. African Journal of Agricultural Research, 13(17), pp.927–933.

Ayele, B., Eshetu B., Betelehem B., Bekele H., Melaku D., 2008. Review of two decades of research on diseases of small cereal crops. In: Abrham Tadesse (eds). Increasing crop production through improved plant protection volume I. Proceedings of 14th annual conference of plant protection society of Ethiopia (PPSE) 19-22 December. 2006 Addis Ababa, Ethiopia 375–416.

Azmeraw, Y., Admassu, B., Abeyo, B., and Bacha, N., 2020. Virulence Spectrum of Puccinia graminis f. sp. tritici in Northwest Ethiopia. Ethiopian Journal of Agricultural Sciences, 30(1), pp.87–97.

Bekele Hundie kotuu, Varkuijl W., Mwangi D., and Tanner G., 2000. Adoption of improved wheat technologies in Adaba and Dodola Woredas of Bale high Land, Ethiopia. Mexico D.F.: CIMMYT.

Belayneh Admassu, Wolfgang Friedtand F. Ordon, 2012. Stem rust seedling resistance genes in Ethiopian Wheat cultivars and breeding lines. African Crop Science Journal, 20(3), pp. 149 – 161.

CSA (Central Statistical Authority). 2019. Report on area and production of major crops (Private peasant Holdings, Meher Season). Addis Ababa, Ethiopia. 1: Pp. 15.

CSA(Central Statistical Authority). 2017. Agricultural sample survey: Report on area and production of major crops (Private peasant Holdings, Meher Season). Addis Ababa, Ethiopia. 1: Pp. 14.

Diriba, Getachew, 2018. Overcoming Agricultural and Food Crisis. Institutional Evolution and the Path to Agricultural Transformation.USA and Addis Ababa.

Edmeades, G., Fischer, R.A., and Byerlee, D., 2010, November. Can we feed the world in 2050? In Proceedings of the New Zealand grassland association. 72, pp. 35–42).

Endale Hailu, Getaneh Woldaeb, Worku Denbel, Wubishet Alemu, Tekelay Abebe, Agengehu Mekonnen, 2015. Distribution of Stem Rust (Puccinia graminis f. sp. tritici) Races in Ethiopia, Plant. Vol. 3, No. 2, 2015, pp. 15–19. doi: 10.11648/j.plant.20150302.11

Enumeration, E.A.S., 2002. Report on the preliminary results of area, production, and yield of temporary crops (Meher season, private peasant holdings). Part I. Central Statistics Authority, Addis Ababa, Ethiopia.

FAO (Food and Agricultural Organization of the United Nations). 2017. Spread of damaging wheat rust continues new races found in Europe, Africa, and Central Asia (b).40pp,

FAO (Food and Agriculture Organization of the United Nations). 2014. FAOSTAT statistical database.

Fetch T.G., Dunsmore K.M., 2004. Physiological specialization of Puccinia graminis on wheat, barley, and oat in Canada in 2001. Canada Journal of plant pathology. 26:1 44–55.

Gomez, K.A. and A.A. Gomez, (1984). Statistical procedures for agricultural research (2 ed.). John Wiley and Sons, New York, 680p.

Haile Selassie, A., Agide, Z., Erkossa, T., Hoekstra, D., Schmitter, P.S., and Langan, S.J., 2016. On-farm smallholder irrigation performance in Ethiopia: from water use efficiency to equity and sustainability.

Hailu, A., Woldeab, G., Dawit, W., and Hailu, E., 2015. distribution of wheat stem rust (Puccinia Graminis F. Sp. Tritici) in West and Southwest Shewa Zones and identification of its physiological races. Advances in Crop Science and Technology, 3(4), p.189.

Hailu Gebre-Mariam, Tanner, D.G., and Mengistu Hulluka, eds. 1991. Wheat Research in Ethiopia: A historical perspective IAR/CIMMYT. Addis Ababa: Ethiopia.

Hei N.B., Tesfaye T., Woldeab G., Hailu E., Hundie B., Kassa D., Yirga F., Anbessa F., Alemu W., Abebe T., Legesse M., Seid A., and Gebrekirstos T., 2018. Distribution and frequency of wheat stem rust races (*Puccinia graminis* f. sp. *tritici*) in Ethiopia. Journal of Agricultural and Crop Research, 6(5), pp.88–96.

Hodson, D. P., Grønbech-Hansen, J., Lassen, P., Alemayehu, Y., Arista, J.,Sonder, K., Kosina, P., Moncada, P., Nazari, K., Park, R. F., Pretorius, Z. A.,Szabo, L. J., Fetch, T., and Jin, Y. 2012. Tracking the wheat rust pathogens. Pages (11–22) in Proc. BGRI Tech.The workshop, Beijing. R. A. McIntosh, ed. Online publication.

Jin Y., Szabo, L. J., Pretorius, Z. A., Singh, R. P., Ward, R., and Fetch, T., Jr. 2008. Detection of virulence to resistance gene Sr24 within race TTKS of *Puccinia graminis* f. sp. tritici. Plant Dis. 92:923–926.

Kumar A., Mishra V.K., Vyas RP. & Singh V. 2011. Heterosis and combining ability analysis in bread wheat (Triticum aestivum L.). Journal Plant Breeding Crop Science. 3: 209–217.

Lemma, A., Woldeab, G., and Semahegn, Y., 2015. Virulence Spectrum of Wheat Stem Rust (Puccinia graminis f. sp. tritici) in the Eastern Showa of Central Ethiopia. Advances in Crop Science and Technology, pp. 1–6.

Leppik, E.E., 1970. Gene centers of plants as sources of disease resistance. Annual Review of Phytopathology, 8(1), pp.323–344.

Mann, M., and Warner, J., 2015. Ethiopian wheat yield and yield gap estimation: a small area integrated data approach. Research for Ethiopia’s Agricultural Policy, Addis Ababa, Ethiopia.

MOA(Ministry of Agriculture). 2020. Irrigation-based wheat production: a transformation from import to export Web: www.eiar.gov.et

Naod Bete-sellassie, Chemeda Fininsa and Ayele Badebo. 2005. The resistance spectra and postulated Sr genes of Ethiopian hexaploid and tetraploid wheat cultivars. In: Proceedings of the 10thAnnual Conference of the Crop Protection Society of Ethiopia, 11-12 August 2005. Ethiopian Agricultural Research Organization (EARO), Addis Ababa. Program and Abstracts. pp. 14–15.

Negatu, W., and Mwangi, W., 1994. An economic analysis of the response of durum wheat to fertilizer: Implications for sustainable durum wheat production in the central highlands of Ethiopia. Developing Sustainable Wheat Production Systems, p.177.

Park R, Fetch T, Hodson D, Jin Y, Nazari K, Prashar M, Pretorius Z., 2011. International surveillance of wheat rust pathogens: progress and challenges. Euphytica 179:109–17.

Park, R.F., 2007. Stem rust of wheat in Australia. Australian Journal of Agricultural Research, 58(6), pp.558–566.

Peterson, R.F., Campbell, A.B., and Hannah, A.E., 1948. A diagrammatic scale for estimating rust intensity on leaves and stems of cereals. Canadian Journal of research, 26(5), pp.496–500.

Roelfs, A.P., and Martens, J.W., 1988. An international system of nomenclature for Puccinia graminis f. sp. tritici. Phytopathology, 78(5), pp.526–533.

Roelfs, A.P., Singh, R.P., and Saari, E.E., 1992. Rust diseases of wheat: Concepts and methods of disease management. CIMMYT, Mexico, DF. Rust diseases of wheat: Concepts and methods of disease management. CIMMYT, Mexico, DF.

Rosegrant, M.W. and Agcaoili, M. (2010), Global Food Demand, Supply, and Price Prospects, International Food Policy Research Institute, Washington, DC.

Sileshi, A., Kibebew, K., & Amanuel, Z. 2015. Temporal and spatial variations in salt-affected soils using Geographical Information System (GIS) and remote sensing at Dubti/Tendaho state farm. [Ph.D.Dissertation thesis] Haramaya University, Ethiopia.

Stakman, E.C., Stewart, D.M., and Loegering, W.Q., 1962. Identification of physiologic races of Puccinia graminis var. tritici. Washington: USDA.

Stubbs, R.I.W., Prescott, J.M., Saari, E.E. and Dubin, H.J., 1986. Cereal disease methodology manual.

Tesemma, T., and Belay, G., 1991. Aspects of Ethiopian tetraploid wheat with emphasis on durum wheat genetics and breeding research.Wheat Research in Ethiopia: A Historical Perspective (Gebre-Mariam, H., Tanner, DG and Hulluka, M., eds), pp.47–71.

Weigand, C., 2011. Wheat import projections towards 2050. US Wheat Associates, USA.

Wuletaw Tadesse, Zewdie Bishaw, and Solomon Assefa. 2018. Wheat production and breeding in Sub-Saharan Africa Challenges and opportunities in the face of climate change. International Journal of Climate Change Strategies and Management. 11(5): 696–715.

Yehizbalem A., B. Admassu, B. Abeyo, and N. Bacha, 2020. Virulence Spectrum of Puccinia graminis f. sp. tritici in Northwest Ethiopia.Ethiop. J. Agric. Sci. 30(1) 87–97.

Zegeye, T., 2001. Adoption of improved bread wheat varieties and inorganic fertilizer by small-scale farmers in Yelmana Densa, and Farta districts of Northwestern Ethiopia. CIMMYT.

